# Electrophysiological profiling of exocytosis during early-stage development of the zebrafish lateral line

**DOI:** 10.1101/2025.11.07.686968

**Authors:** Jiali Wang, Erdem Karatekin, David Zenisek

## Abstract

Hair cells of the zebrafish lateral line have proven to be a good model for studying hair cell function in a system that is easily genetically manipulated, rapidly develops and is experimentally accessible. However, characterization of potential developmental changes, and possible differences along lateral line position are lacking. Here, we used *in vivo* patch clamp to investigate the electrophysiological and exocytic properties of neuromast hair cells over early development across body location. Long depolarizations led to steady increases in membrane capacitance, presumably due to exocytosis of vesicles localized to ribbon synapses. The magnitude and kinetics of capacitance changes did not vary significantly across the L1 to L6 position of neuromasts along lateral line, but the magnitudes were found to be significantly smaller in hair cells found in the tail region across all developmental time points. For each region, we found no significant changes in capacitance responses between 3 and 7 days after fertilization. Hair cell capacitance responses were greatly reduced in animals injected with CRISPR/Cas9 with gRNAs targeted to otoferlin b. These results confirm the essential role of otoferlin b in neuromast hair cell function, and they establish the fidelity of CRISPR/Cas9 to rapidly mediate genetic removal of critical genes to study their impact on synaptic release.

## INTRODUCTION

Zebrafish hair cells play an important role in both the auditory and vestibular systems. Lateral line hair cells—located in mechanosensory organs called neuromasts along the body, serve as a functional analog to mammalian hair cells. These neuromasts detect water flow and are essential for survival behaviors such as prey detection, mating, and predator avoidance (1). Due to their structural and functional similarities to mammalian hair cells, the zebrafish lateral line system provides an excellent model for studying synaptic transmission, hair cell regeneration, and hearing-related mechanisms (2).

A key feature of hair cells is the synaptic ribbon, an electron-dense structure at the presynaptic active zone that tethers synaptic vesicles (SVs). These ribbons facilitate rapid exocytosis and signal transmission upon mechanical stimulation. Mature hair cells in zebrafish contain approximately 130 vesicles per ribbon and 2-5 ribbons per hair cell (3-6).

The lateral line develops early in zebrafish embryogenesis. By 48 hours post-fertilization (hpf), a migrating primordium places clusters of cells, each forming a neuromast (L1–L6), with additional terminal neuromasts developing at the tail tip. Intercalary neuromasts later arise from proliferating inter-neuromast cells (7-9). The lateral line develops rapidly in zebrafish, with hair cells emerging at 48hpf and becoming capable of responding to sensory stimuli within 48 hpf (9-17), and reaching maturity by 5 dpf (18,19). However, functional heterogeneity among neuromasts remains unclear, a comprehensive electrophysiological analysis of synaptic function across neuromasts is lacking, the timing of electrophysiological maturation has not been fully characterized.

In this study, we employed *in vivo* whole-cell electrophysiology to investigate calcium currents and exocytosis (20,21) in lateral line hair cells during early development (3–7 days post-fertilization, dpf) and across different neuromasts. We additionally tested the role of otoferlin, a critical protein in hair cell synaptic vesicle release, using CRISPR-injections. We found significant impairments in exocytosis in *otof* mutants that did not vary across developmental time. Thus, unlike in mammalian inner hair cells where Otoferlin expression gradually replaces that of Synaptotagmin-1 within the initial 14 days after birth (22), in zebrafish, otof b expression is present around the time of synapse formation, and is functionally non-redundant. Our results further establish that CRISPR/Cas9 injections can be used in acute studies to monitor effects of protein deletion at the single cell level. Interestingly, we identified variations in function between neuromasts, particularly those at the tail in comparison to the body. Finally, we combined immunostaining of pre- and post-synaptic markers and analysis of otoferlin expression to assess hair cell functionality. We found that during 3-7 dpf, hair cell numbers increase, and synapses mature, but release properties of individual hair cells change little. These results highlight functional diversity among neuromasts and underscore the zebrafish lateral line as a valuable model for synaptic physiology and hearing research.

## RESULTS

### Electrophysiological characterization of zebrafish neuromast hair cells across body locations during development

To study hair cells of neuromasts, we used the patch clamp technique, as described previously (20). The procedure involves perforating the neuromast periphery with a large-bore pipette to allow access of a patch pipette to the hair cell membrane. After achieving whole-cell configuration, the membrane capacitance C^m^, reflecting cell membrane area (23,24), was continuously recorded. We used a high-frequency dual-sine approach that allows to track and record capacitance despite the presence of membrane currents (3,21). For all recordings, we used a Cs^+^-based internal solution to block K^+^ channels. We found no evidence of transient Na^+^ currents in our experiments, which suggests a lack of voltage-gated Na^+^ channels. This contrasts with developing mouse hair cells, which do exhibit these channels and action potentials in immature hair cell (25). Since hair cells are typically known to lack other inward conductances (20,26-28), the remaining current was presumed to reflect current from the L-type Ca^2+^ channels that drive neurotransmitter release (3,20,29,30).

The posterior lateral line forms between 19 and 25 hpf (31-33). Zebrafish possess neuromasts distributed along the anteroposterior body axis, defined as L1-L6 covering the surface of the lateral line. Neuromasts contain hair cells that develop in pairs, with their stereocilia facing opposite directions, enabling detection of both forward and backward water flow (10,34-37). Additionally, three neuromasts develop outside the lateral line and are referred to as “secondary neuromasts.” These neuromasts are sensitive to water movements along perpendicular axes (37). The neuromasts aligned along the anteroposterior body axis are termed primary neuromasts and are labeled L1 to L6 (10,37-39). The secondary neuromasts, which are perpendicular, are labeled LII1 to LII3 (37,40). In addition to these neuromasts, two neuromasts are found on the tail. By 48 hpf, the embryonic lateral line pattern is fully established, but hair cells continue to develop over the next several days (18,19,41). However, how electrophysiological properties vary across different neuromast locations and across developmental time is not fully understood. To address this question, we recorded from zebrafish between 3 and 7 dpf in neuromasts across the body and tail and subjected hair cells to 5 s depolarizations to −10 mV while monitoring C_m_ as an index of exocytosis (Fig. 1A). Average capacitance traces of hair cells at the L1 or L3 positions at different developmental timepoints are shown in Fig. 1B-D. Capacitance rises with depolarization in an exponentially slowing pattern as previously described (3,20). This waveform persisted regardless of location and time during development (Figure 1B-D).

**Fig. 1.**
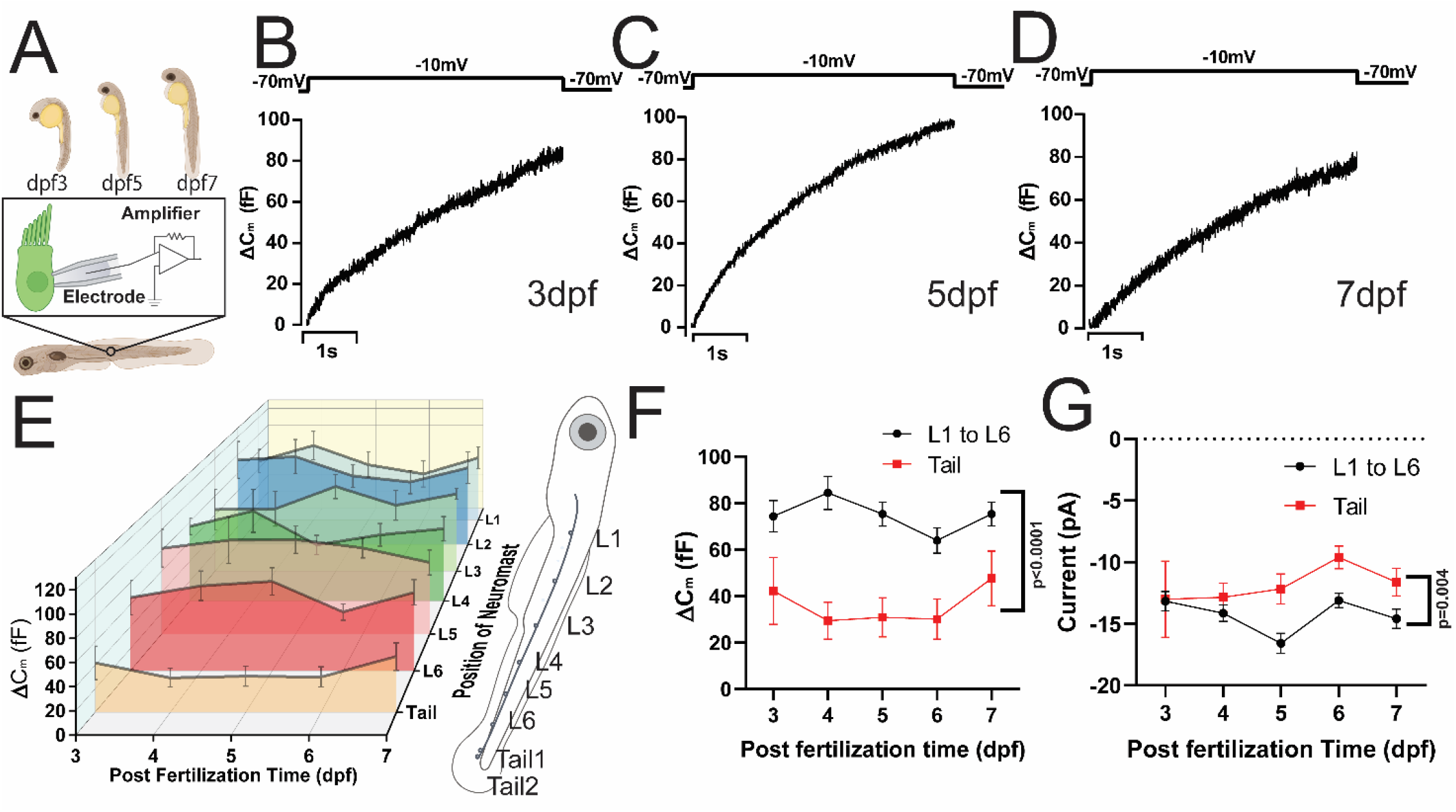
Exocytosis in neuromast hair cells across locations and developmental time. (A) Diagram of zebrafish development from 3 to 7 days post-fertilization (dpf) and illustration of whole-cell recordings. (B-D). Capacitance responses to 5 s depolarizations to −10 mV in zebrafish neuromasts from 3 to 7 dpf taken from L1, L3, and L3 respectively. (E) Changes in capacitance (Δ*C*_*m*_) in response to 5 s depolarizations at different developmental times and neuromast positions. The neuromasts at the tail consistently showed smaller changes, we observed similar level of Δ*C*_*m*_ from L1 to L6 and combined these together as one group as plotted in (F-G). (F) The same data set as in E, but comparing capacitance changes averaged across neuromasts in L1 to L6 to those at the tail at different development points (t-test, p<0.0001). (G) Same as in F, but y-axis indicated the negative peak current from I-V ramp recording during depolarization from −80mV to +40mV (t-test, p= 0.004).

We systematically recorded from L1 to L6 and tail 1 and tail 2 neuromasts from 3 to 7 dpf. A 3-dimensional plot summarizes hair cell capacitance responses to 5 s depolarizations as a function of development time (dpf) and neuromast location (Figure 1E). Quantification of capacitance changes showed little variation over developmental time whether in the body or the tail regions. For example, the mean ΔC_m_ for L3 hair cells was 65.9 ± 21.4 fF at 3 dpf, 66.5 ± 18.4 fF at 4 dpf, and 82.3 ± 5.7 fF at 7 dpf (Fig. 1E). Statistical analysis confirmed no significant differences across the different time points or either analysis on L1-6, or tail locations (p>0.05; ANOVA test).

We also found that the average capacitance response was largely consistent across neuromasts L1 to L6 at all developmental stages. For example, following a 5 s depolarization at 7 dpf, the mean capacitance response ΔC_m_ was 72.8 ± 12.6 fF for L1, 85.5±17.6 for L2, 82.3 ± 5.7 fF for L3, 73.1±12.3 for L4, 68.7±11.2 for L5, and 71.1 ± 11.5 fF for L6, with no statistical difference among the measurements (p>0.05 in ANOVA test).

Recordings from hair cells located on the tail neuromasts showed little variation over time as mentioned, but their capacitance responses were consistently smaller than those in the L1-L6 neuromasts. Since responses varied little from L1 to L6, we grouped responses from these neuromasts for subsequent analysis, as shown Fig. 1F and 1G. Capacitance responses were significantly smaller in tail hair cells than those in the body across all developmental time points measured (p<0.0001, Student’s t-test). We next compared calcium currents elicited by 5 s depolarizations over developmental time beginning at 3 dpf and continuing until 7 dpf. Negative peak currents were similar at 3 and 4 dpf (p=0.48 for 3dpf, p=0.18 for 4dpf) for locations at the tail compared to L1-L6, but had smaller amplitude at later time points (Fig. 1G). Thus, the lower magnitude of Ca^2+^ currents on the tail neuromasts may contribute to their smaller capacitance responses compared to L1-L6 neuromasts for 5-7 dpf.

### Minor alterations in hair cell synapses in early development

Since we found that exocytosis was similar between 3 dpf and 7 dpf for a given neuromast, but changed based on location, we next investigated whether hair cell and synapse numbers also changed across location and time. To do so, we immunostained for the synaptic ribbon protein, Ribeye, and post-synaptic marker pan-MAGUK, as done previously by other investigators (3,5,42) (Fig. 2A-D). Ribeye is a gene product arising from an alternate start site to the transcriptional co-repressor CtBP2 (43) and thus can be immuno-stained using antibodies that recognize CtBP2. We defined assembled ribbon synapses as ones that contained both pre-synaptic ribbon (CtBP2) and post-synaptic (pan-MAGUK) markers in close proximity and immature or ribbon-precursors (4,44) as ribeye spots that lacked nearby (< 0.5μm) post-synaptic markers (45). The number of Ribeye spots, pan-MAGUK spots and colocalized pan-MAGUK/Ribeye spots increased from day 3 to day 7. The number of co-localized markers was nearly identical in both the body and tail at 3 dpf (body 5.9±0.4; tail 6.0 ±0.6, p=0.279) and 7 dpf (body 25.1±0.9; tail 24.1 ± 1.3, p=0.06), so we combined the data for presentation in Figure 2E.

**Fig. 2.**
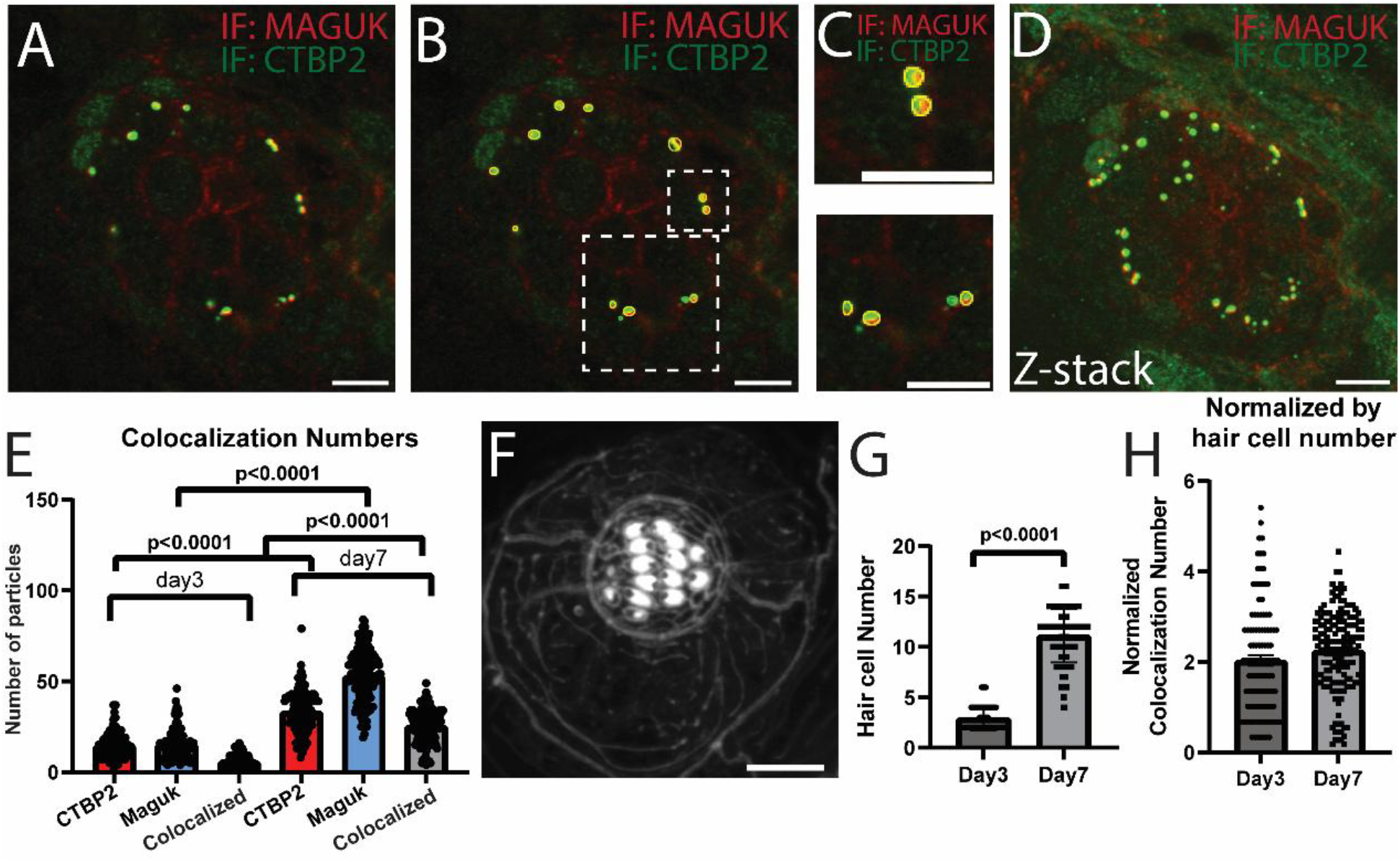
Synapse Formation from Day 3 to Day 7. (A) Immunostaining of CTBP2 (Green) and MAGUK (Red) in 3 dpf zebrafish. (B) The same image as in A, but with assembled synapses marked by a yellow outline. Assembled synapses are defined by pre- and post-synaptic markers, CTBP2 and MAGUK, respectively, co-localized within 0.5 µm. Post-synaptic markers (red) are lacking near some presynaptic markers (green), indicating some synapses are yet to be assembled. (C) Magnified view of regions in B. (D) Z-projection from the same neuromast shown in (A-C), Z-stack view of pre-(CTBP2, Green) and post-synaptic markers (MAGUK, Red) taken from 48 sections over 12.69 mm total distance. (E) The number of pre- and post-synaptic marker spots and their colocalization statistics, calculated from z-stack images as in D, using ImageJ (72), shown for 3 and 7dpf animals. This analysis included 112 neuromasts at 3 dpf and 164 neuromasts at 7 dpf. (F) A confocal microscopy z-stack projection of a neuromast, viewed from the top. The neuromast was from 7 dpf larva, at position L1. Hair cell numbers were evaluated from cilia staining by fluorescent phalloidin. (G) The number of hair cells per neuromast. There were 3.0 ± 0.2 and 11.6 ± 0.2 hair cells per neuromast at 3 and 7 dpf, respectively. (H) The number of colocalized CTBP2/pan-MAGUK spots per neuromast, normalized to the number of hair cells, The average number of colocalized spots per hair cell was 2.03±0.12 at 3 dpf and 2.25±0.1 at 7 dpf. (t-test, p=0.1).

Hair cell numbers also increase over this period of development (46-48), which may partially or completely account for the change in ribbon numbers. To account for hair cell proliferation, we normalized the number of co-localized spots to the number of cells in each neuromast. To identify hair cells, we used phalloidin to visualize hair cell bundles (10) at the apical tip of hair cells and counted the number of hair bundles in each neuromast at 3 dpf and 7 dpf (Fig. 2F). The number of hair cells per neuromast was consistent across the two locations at day 3 (body 5.9 ± 0.4, tail 6.0 ±0.6) and day 7 (body 25.1 ± 0.9, tail 24.1 ± 1.3). The combined data is presented in Fig. 2G. When normalized to cell count at each developmental stage, there was no difference in the number of spots containing adjacent pre- and postsynaptic markers, as shown in Fig. 2H (2.03±0.12 at 3 dpf; 2.25±0.1 at 7 dpf, t-test, p=0.1).

We next examined recruitment of Ribeye to synapses across location and development time. For this, we used immunostaining of Ribeye and MAGUK as above, but focused on the fraction of Ribeye spots (marked by anti-CtBP2 antibodies) that co-localized with spots marked by the post-synaptic Maguk (Fig. 3A-G). There was no difference between the body and tail regions at a given dpf. However, the percentage of Ribeye spots colocalized with post-synaptic Maguk spots nearly doubled from ∼40% at 3 dpf to ∼75% at 7 dpf for both regions (Fig. 3 H,I). Combined with results above, this implies that the number of extra-synaptic Ribeye spots per cell must have decreased from 3 to 7 dpf, reflecting recruitment of Ribeye to synapses. Our results also indicate that while the degree of co-localization of pre-to post-synaptic markers changed dramatically over this period, the synapse number alone cannot account for the differences in capacitance response magnitudes in the tail region relative to the body.

**Fig. 3.**
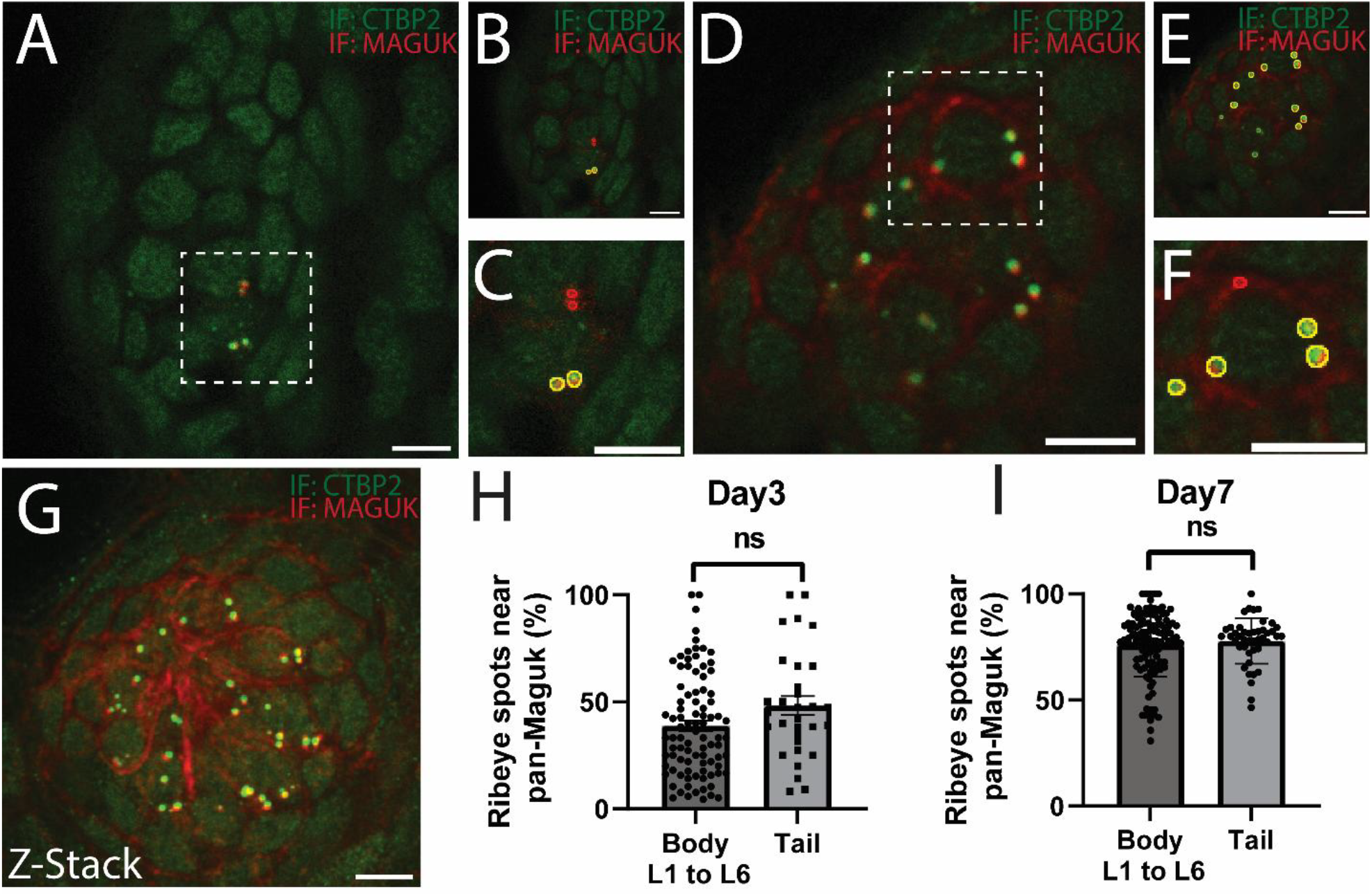
Ribbon synapse development in tail and body. (A-F) Immunostaining of ribbon markers CTBP2 (green) and postsynaptic marker pan-MAGUK (red) in day 3 (A-C) and day 7 (D-F) animals. Examples of colocalized CTBP2 and pan-MAGUK are outlined in yellow and non-colocalized spots are outlined in red (pan-MAGUK) or green (CTBP2). B,C and E,F are magnified regions from A and D, respectively. (G) Z-stack with maximum intensity projection of a day 7 neuromast across 54 confocal slices covering 12.19 μm. (H-I) Synapse development, assessed based on the percentage of pre-synaptic spots that co-localized to post-synaptic spots for 3 dpf (H) and 7 dpf (I) animals. The number of CtBP/ribeye spots (presynaptic, Green) and pan-MAGUK spots (postsynaptic, red) per neuromast were counted at L1-L6 and tail locations. A total of 141 hair cells for 3 dpf and 168 hair cells 7 dpf were analyzed. No significant difference between tail and body were found at 3 dpf (p =0.07, Student’s t-test) or 7 dpf (p= 0.7).

Surprisingly, these results suggest that both synapse number and exocytic capacity remain unchanged during early development, despite the poor localization of Ribeye to pan-MAGUK at early time points and other well-documented changes in properties that hair cells undergo over this period (29,49,50). It is worth noting that genetic elimination of ribbon synapses does not cause a reduction in capacitance responses in either zebrafish (3) or mammalian hair cells (51,52), consistent with our findings that hair cells that have not finished recruiting Ribeye to assembled ribbon synapses can support normal amounts of exocytosis.

### Otoferlin expression in lateral line hair cells

As an additional test for visualizing hair cells and to assess developmental changes to proteins involved in synaptic transmission, we investigated otoferlin expression. Otoferlin is a key protein required for hair cell exocytosis, which, when mutated, leads to inherited forms of human deafness. To date, 296 mutations and deletions in the otoferlin gene (*Otof*) have been identified, resulting in a spectrum of auditory impairments ranging from mild hearing loss to profound deafness (53,54), and otoferlin-deficient mice (*Otof*−/−) exhibit profound deafness (55). In these mice, exocytosis in inner hair cells and type I vestibular hair cells are nearly abolished despite normal ribbon synapse morphology and Ca^2+^ currents (55). These findings underscore otoferlin’s essential role in synaptic vesicle exocytosis and suggest it may serve as the primary Ca^2+^ sensor triggering membrane fusion at the IHC ribbon synapse (56).

During mammalian development, inner hair cells undergo a developmental switch that shifts from a mode of neurotransmitter release that is Synaptotagmin-1 dependent to one that requires Otoferlin (22). Here, zebrafish were collected at 3 dpf and 7 dpf to be fixed and stained for Otoferlin. Upon imaging, neuromasts were readily apparent as the most intensely stained structures within the zebrafish, consistent with high expression levels of otoferlin, Fig. 4A. Our results indicated no significant differences in intensity between day 3 and day 7 at each location (Fig. 4B,C). Notably, otoferlin staining revealed lower intensity in hair cells located on the tail compared to those on the body, possibly indicating a regional variation in otoferlin expression and consistent with lesser vesicle trafficking in the tail region (Suppl.Fig1).

**Fig. 4.**
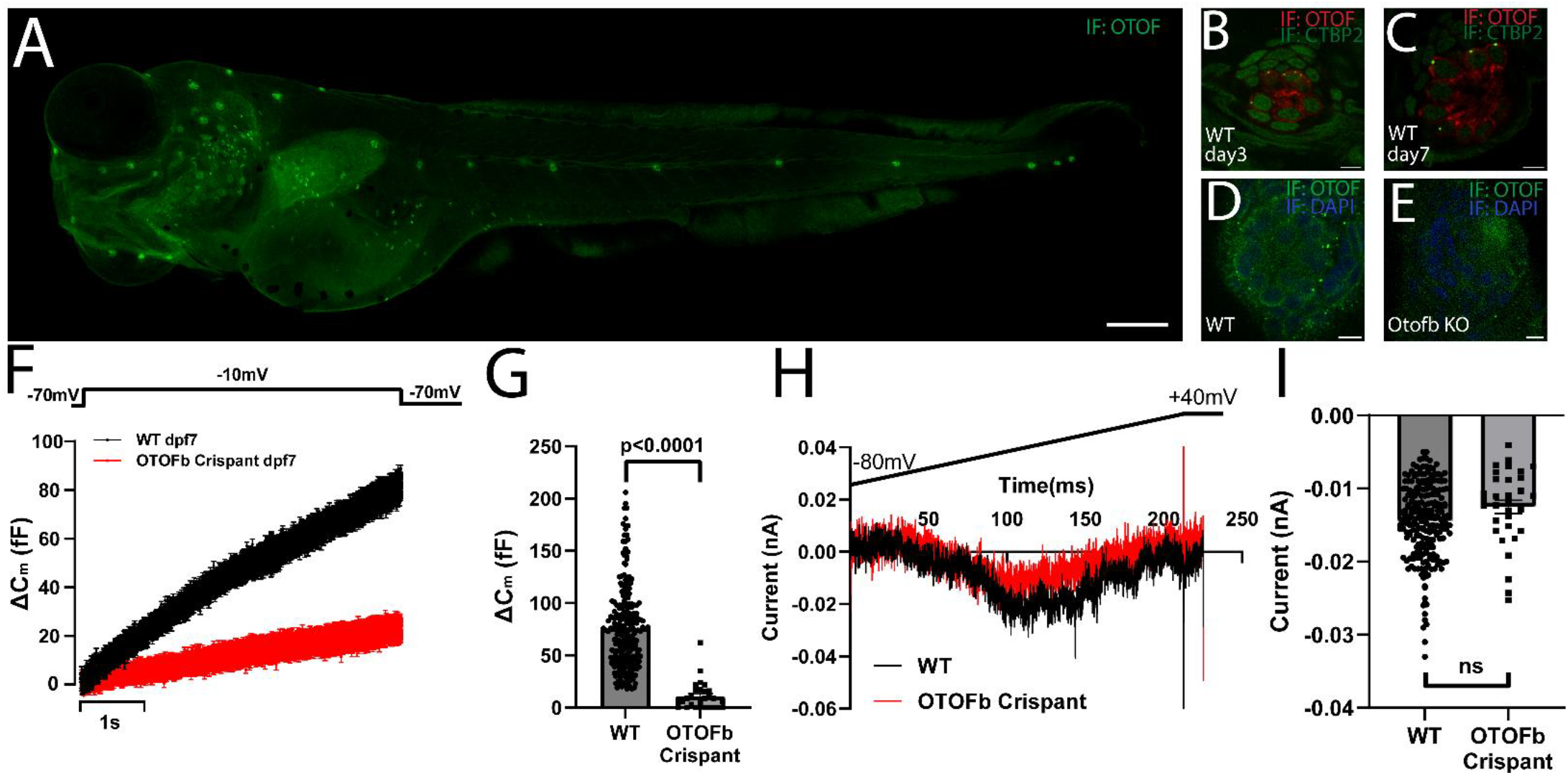
Otoferlin b-mediated exocytosis in zebrafish neuromast hair cells over development. (A) Immunostaining for otoferlin (Green) in whole fish at 6 dpf. (B) Otoferlin (red) and CTBP2 (green) staining of a 3 dpf neuromast. (C) Same, for 7 dpf. (D) Immunostaining of Otoferlin (green) in a neuromast from a WT animal. DAPI (blue) labels nuclei. (E) Same as in D, but for otoferlin b crispants. (F) Whole-cell capacitance measurements of 7 dpf WT (black) and Otoferlin crispant (red) zebrafish in response to 5s depolarizations from −70mV to −10mV. The traces are averages from 6 recordings each at position L3. (G) Average ΔC_m_ of WT (n = 228) and *otofb* gRNA-injected embryos at dpf 5-7 (n = 20) neuromast hair cells in response to 5 s depolarizations to −10 mV. The capacitance change was 76.4 ± 2.6 fF (mean ±SEM) for WT hair cells and 15.4 ± 3.0 fF for *otofb* mutants. (H) Typical currents in response to a voltage ramp for WT (black) and otoferlin b crispant animals (red). (I) Average calcium currents are shown, for WT (n=231) and *Otofb* (n=32) 7 dpf animals injected with gRNA as embryos, in response to 5 s depolarizations to −10 mV. The inward calcium peak current was −14.4 ± 0.3 pA for WT hair cells and −12.5 ± 0.9 pA for mutants.

### Otoferlin b plays an important role in zebrafish lateral line hair cells

Zebrafish have two Otoferlin genes, designated *otoferlin a* and *otoferlin b. otoferlin b* (*otofb*) has been shown to be the dominant isoform in neuromasts (57). To investigate the role of Otoferlin b in hair cell exocytosis, CRISPR-Cas9 protein was co-injected with two gRNAs targeting exon 39 and 40 of *otofb* at the one-cell stage to generate *otofb* crispants. A subset of crispants were collected for immunostaining to assess Otof protein expression and CRISPR efficiency. As expected, CRISPR-Cas9 injected larvae displayed far less Otoferlin intensity as shown in Fig. 4 (D–E).

Additionally, crispants from days 3 to 7 were used for *in vivo* membrane capacitance recordings in neuromast hair cells. *Otofb* crispants showed a dramatic reduction in capacitance response to 5 s depolarizations, as shown in Fig. 4F. On average, WT hair cells from L1 to L6 exhibited an increase of 76.4 ± 2.6 fF during depolarization, whereas in *otofb* crispants, the response was significantly reduced to 15.4 ± 3.0 fF (Fig. 4G, t-test, p<0.0001), suggesting the absence of an otoferlin-independent phase of exocytosis during this developmental time period.

We also assessed calcium currents during whole-cell voltage clamp experiments. To do so, cells were recorded with a solution designed to block most K^+^ currents (see methods). An inward Ca^2+^ current was observed during both I-V ramp and I-V step protocols, with the membrane potential stepped from −80 mV to +40 mV. The average negative Ca^2+^ peak current in WT hair cells was −14.4± 0.3 pA. In *otofb* crispants, the Ca^2+^ influx trended smaller by approximately 20%, with an average current of −12.5± 0.9 pA (Fig. 4I), though the diference did not reach statistical significance (t-test, p = 0.075).

## DISCUSSION

### Hair Cell exocytosis development from dpf 3 to dpf 7

Hair cells located in the neuromasts of the lateral line system are known to play a critical role in detecting water flow and predators (1,58-61). Deflections of stereocilia lead to depolarization and release of glutamate via exocytosis. Here, we examined how these properties change along the axis of zebrafish and during development, by collecting electrophysiological data from 3 to 7 dpf at different neuromast locations. During this period, hair cells undergo developmental changes (18,19,29,49,50), so changes in mechano-transduction properties are expected. Indeed, previous literature indicates that at 3 dpf, neuromasts contain immature hair cells with reduced mechano-transduction capabilities. Specifically, hair cells from 3 dpf animals display reduced afferent firing rates compared to 7 dpf animals (46). However, unexpectedly, we observed no significant differences in exocytosis, probed by capacitance recordings. A possible explanation for this is that the exocytosis capacity of neuromast hair cells develops before the maturation of synaptic connections with afferent neurons. Using pre- and post-synaptic markers to probe synapse development, we found that synaptic connections undergo significant development between day 3 and day 7, as the percentage of ribeye spots that localize near post-synaptic markers increases dramatically over this period, consistent with previous reports (29). Interestingly, the number of synapses per cell remains roughly the same at about two per synapse through 3 to 7 dpf, but the number of extra-synaptic Ribeye spots decreases, reflecting the recruitment of Ribeye to synapses. Our results suggest that the ability to support exocytosis precedes the development of mechano-transduction and full synapse assembly.

The ability of presumably immature ribbons to support high release rates is consistent wih previous studies on ribbon synapses lacking ribbons. While synaptic ribbons are a hallmark feature of auditory, vestibular, and visual systems, that function to tether synaptic vesicles near the presynaptic release sites, they may be dispensable for achieving maximal release rates from some ribbon synapses. Previous studies have demonstrated that cells with ribbon synapses exhibit sustained exocytosis in response to prolonged stimuli (27,62-66). However, while fish (3) and mice (51,52,67) lacking Ribeye also lack ribbons, they are still capable of supporting high exocytic rates.

### Zebrafish lateral line as a model for studying exocytosis

In our study, the zebrafish larva presents a versatile model for the study of exocytosis. Its unique advantages, particularly the large numbers of offspring, external development, direct access for in vivo electrophysiology, rapid development and accessible genetic tools, allow for rapid molecular manipulation and functional studies. The rapid generation of CRISPR/Cas9-based crispants within a matter of days, as demonstrated in this study, permits the efficient evaluation of genetic contributions to exocytotic mechanisms. This combination of physiological and genetic tractability establishes the zebrafish larva as an ideal platform for accelerating the discovery of gene functions regulating vesicular release.

Despite these advantages, rapid developmental changes in the expression of proteins involved in vesicular trafficking may occur and confound interpretation. A protein critical for exocytosis in hair cells is Otoferlin (55,68,69), so we tested its expression in early development in neuromasts. Previous reports in inner hair cells in mammals have demonstrated a developmental (70) switch from a Synaptotagmin-dependent process early in development to an otoferlin-requiring process as hearing onsets after birth (22). Here, we found that reduction of Otoferlin dramatically reduced the exocytic response in both 3 dpf and 7 dpf animals to a similar extent. This suggests that neuromast hair cells in zebrafish, unlike mammalian inner hair cells, either do not pass through an otoferlin-independent stage or do so prior to 3 dpf. This makes neuromast hair cells an attractive model to study Otoferlin function.

### Distinct functions and patterns of tail neuromasts

Our recordings revealed differences in the capacitance responses to depolarization for hair cells localized to the neuromasts located on the tail in comparison to those on the body: exocytosis was reduced by ∼40% during depolarization in the tail region. This characteristic may reflect distinct biological roles and adaptations to the different mechanical sensory environments of each neuromast region. Specifically, neuromasts L1-L6 are strategically positioned to detect water flow from forward and backward directions, a function critical for spatial orientation and predator detection (1). Exocytosis is proportional to stereocilia deflection, hair cells particularly those near the head and trunk, are often more exposed to subtle environmental changes than the tail. In these regions, smaller stereocilia deflections may couple with higher exocytotic sensitivity to ensure reliable signaling. In contrast, the tail’s primary function is thrust generation, resulting in proportionally larger stereocilia deflections. The reduced exocytotic response we observed in tail hair cells could prevent signal saturation, effectively “turning down the volume” to maintain the range of signaling for tail-generated stimuli. Nevertheless, our data show that synaptic density per hair cell remains consistent. These results emphasize that studies of electrophysiological and synaptic properties of zebrafish hair cells should make note of these differences and account for them when interpreting their results. Furthermore, these results imply that structural and/or molecular differences in the tail region may differ from those in the body, including differences in calcium currents and Otoferlin expression. Further differences and mechanisms remain to be elucidated.

## ACKNOWLEDGEMENTS

We thank all members of the Zenisek and Karatekin labs for critical discussions, especially Jie Zhu and Ane Landajuela. We are grateful to Prof. Genglin Li (Eye & ENT Hospital of Fudan University) for demonstrating hair cell recording, and to Ane Landajuela for validating the Otoferlin antibody. This work was supported by the National Institutes of Health (NIH) grants R01NS122388 (to EK and DZ) and R01DC019057 (to DZ and EK). The content is solely the responsibility of the authors and does not necessarily represent the official views of the NIH.

## Author Contributions

J.W. designed the research, performed electrophysiology, immunostaining, genetic perturbations and zebrafish embryo injections, analyzed the data, and wrote the paper. E.K. and D.Z. designed the research and assisted with manuscript preparation.

## MATERIALS AND METHODS

### Zebrafish Husbandry

Zebrafish were kept in accordance with the Yale University Animal Care and Use Committee guidelines. The *otofb* crispant has been generated by Crispr-Cas9 gene editing method. Crispr-Cas9 protein (Integrated DNA Technologies, Inc) as well as gRNA were injected into WT zebrafish embryo at the 1-cell stage. The gRNAs targeting *otofb* were selected from CHOPCHOP and produced by Integrated DNA Technologies, Inc. The gRNA used were TCGGCCTGCAGGTAGTCGAAAGG and GACCTGAATCGATTTCCACGAGG target exon 39 and 40 of otoferlin b. Genotyping was used to confirm injection efficiency by using standard PCR followed by a Sanger sequencing. *otofb* crispant were compared with wild type as stated in the text.

### Zebrafish hair cell electrophysiology

Zebrafish of either sex ranging in age from 3 to 7 days postfertilization (dpf) were anesthetized in Tricaine and mounted by stainless steel pins (Fine Science Tools: 26002-10) in a recording chamber. An upright Olympus microscope was used for viewing, and recordings were made with an Axon 200B amplifier with an Axon DD1322 digitizer, following previous publication (20). All recordings were made with jClamp software (Scisoft). Cells were held at −60 mV. The extracellular solution contained (in mM): 125 NaCl, 1.0 KCl, 2.2 MgCl_2_, 2.8 CaCl2, 10 HEPES, 6 d-glucose, 285 mOsm, pH 7.6. The pipette was composed of (in mM): 90 CsCl, 20 TEA, 5 Na2ATP, 3.5 MgCl_2_, 10 HEPES, 1 EGTA, 260 mOsm, pH 7.2. Pipette resistance was typically 4-5 MΩ with Cs pipette solutions. Patch pipettes were pulled from 1.5-mm outside diameter thick-walled borosilicate glass (World Precision Instruments,TW150F-3) and coated with wax (Sybron Dental Specialties, #000625). Capacitance measurements were made with a dual sine admittance technique (21). Ca^2+^ currents were measured by a rapid 500ms I-V ramp from −80mV to +40mV, or I-V jumps from −80mV to +40mV. As mentioned in results section, no Na^+^ currents were observed and current recorded represent the L-type Ca^2+^ channels activities (3, 20, 29, 30). Recordings were made at room temperature. Data is reported as mean ± SEM. Student’s t-tests or ANOVA followed by post hoc Bonferroni test to determine whether differences between group means were statistically significant.

### Immuno-fluorescence

Zebrafish hair cell staining was performed as previously published (71), 3-7 dpf zebrafish larvae were anaesthetized and fixed in 4% paraformaldehyde in PBS for 6 hours. Next, fish were transferred to blocking buffer consisting of 2% bovine serum albumin, 3% normal goat serum, and 0.1% Triton X-100 in PBS, followed by incubating overnight with primary antibodies diluted in blocking buffer at 4°C. The samples were washed three times with PBS. Next, they were incubated with secondary antibodies for 4 hours at room temperature, followed by additional washes. Finally, the samples were mounted and prepared for imaging.

The primary antibodies used for Anti-Otoferlin, selected from Picoband (PB9616, Boster Biological Technology, USA) and pan-MAGUK (Neuromab) were diluted in 1:1000 and 1:500, respectively. CtBP Antibody (B-3) was diluted in 1:1000 (sc-55502, Santa Cruz Biotechnology Inc.). Both the pan-MAGUK and CtBP antibodies have been used extensively in zebrafish for localizing ribbons and afferent dendrites (3,5,29,42,49,50). The secondary antibodies (Alexa Fluor 488/643 goat anti mouse/rabbit IgG(H+L), Molecular Probes, USA) were diluted 1:200. The secondary antibodies (Alexa Fluor 647 goat anti rabbit IgG1, Molecular Probes, USA) Alexa Fluor 488 goat anti rabbit IgG2a, Molecular Probes, USA) were diluted 1:200.

Confocal images were acquired with a Zeiss (Germany) LSM 950 laser-scanning confocal using Plan-Apochromat 63x/1.40 Oil DIC M27 objective and 1x or 2x digital zoom. Digital images were processed by Image J and Adobe illustrator software. For quantifying image intensity of otoferlin staining, the following parameters were used for all experiments: detector gain at 700V for 488 mm wavelength, 1% intensity, 650V for 405mm wavelength, 2% intensity. Image capture size was set to 1024 x1024.

### Staining analysis

Quantitative image analysis was performed on raw images using Image J (Fiji) software. For analysis, z-stacks collapsed into single z-projections with max intensity. Analysis of colocalization was carried out using spots colocalization plugin. Estimated particle size and intensity threshold were set individually to maintain consistency. An entire neuromast was set into ROI, particles were set as oval to be detected.

To quantitatively measure immunolabel density, we used identical imaging settings to compare the 3dpf to 7 dpf. ROI of auto detection number of CtBP, or MAGUK were evaluated as well. Intensity and number of ROI information were extracted from imageJ and organized in Excel and plotted in Origin, GraphPad or Excel. Student’s t-test was used to determine whether differences between group means were statistically significant.

## Suppl. Figs

**Suppl. Fig.1.**
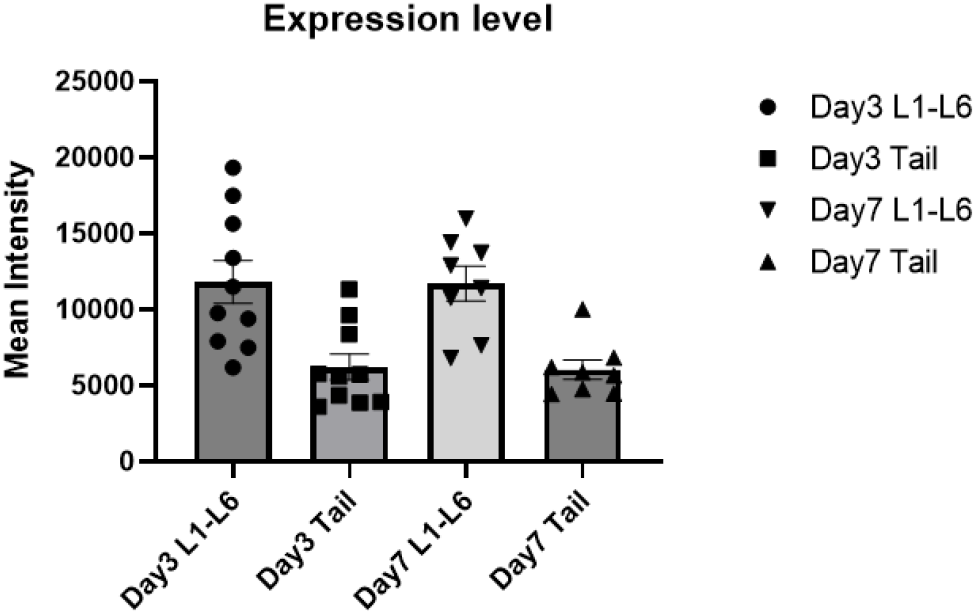
Otoferlin expression is relatively lower in tail neuromasts. Otoferlin expression was quantified via immunofluorescence staining in hair cells from the L1-L6 (body) and tail neuromasts at two developmental time points (day 3 and day 7). At day 3, we analyzed 59 neuromasts from the body and 17 from the tail across 10 larvae. By day 7, we quantified expression in 46 neuromasts from the body and 16 from the tail across 9 larvae. Analysis of the mean fluorescence intensity for each neuromasts revealed consistently higher otoferlin expression in hair cells of the L1-L6 region compared to tail hair cells at both day 3 and day 7.

